# Defined microbiota transplant restores Th17/RORγt^+^ regulatory T cell balance in mice colonized with inflammatory bowel disease microbiotas

**DOI:** 10.1101/844662

**Authors:** Graham J. Britton, Eduardo J. Contijoch, Matthew P. Spindler, Varun Aggarwala, Gerold Bongers, Lani San Mateo, Andrew Baltus, Anuk Das, Dirk Gevers, Thomas J. Borody, Nadeem O. Kaakoush, Michael A. Kamm, Hazel Mitchell, Sudarshan Paramsothy, Jose C. Clemente, Jean-Frederic Colombel, Marla C. Dubinsky, Ari Grinspan, Jeremiah J. Faith

**Affiliations:** The Precision Immunology Institute, Icahn School of Medicine at Mount Sinai; Icahn Institute for Data Science and Genomic Technology, Icahn School of Medicine at Mount Sinai; Department of Oncological Sciences, Icahn School of Medicine at Mount Sinai; Janssen Research & Development LLC; Janssen Human Microbiome Institute, Janssen Research and Development, LLC; Centre for Digestive Diseases, Sydney, Australia; School of Medical Sciences, University of New South Wales, Sydney, Australia; Department of Gastroenterology, St Vincent’s Hospital, Melbourne, Australia; Nambour General Hospital, Nambour, Australia; Bankstown-Lidcombe Hospital, Sydney, Australia; The Dr. Henry D Janowitz Division of Gastroenterology, Icahn School of Medicine at Mount Sinai

## Abstract

The building evidence for the contribution of microbiota to human disease has spurred an effort to develop therapies that target the gut microbiota. This is particularly evident in inflammatory bowel diseases, where clinical trials of fecal microbiota transplant have shown some efficacy. To aid the development of novel microbiota-targeted therapies and to better understand the biology underpinning such treatments, we have used gnotobiotic mice to model microbiota manipulations in the context of microbiotas from humans with inflammatory bowel disease. Mice colonized with IBD donor-derived microbiotas exhibit a stereotypical set of phenotypes, characterized by abundant mucosal Th17 cells and a deficit in the tolerogenic RORγt^+^ Treg cell subset. Transplanting healthy donor-derived microbiota into mice colonized with human IBD microbiotas lead to induction of RORγt^+^ Treg cells, which was associated with an increase in the density of the microbiotas following transplant. Microbiota transplant reduced gut Th17 cells in mice colonized with a microbiota from a donor with Crohn’s disease. By culturing strains from this microbiota and screening them in vivo, we identified a specific strain that potently induces Th17 cells. Microbiota transplants reduced the relative abundance of this strain in the gut microbiota, correlated with a reduction in Th17 cells.

## Introduction

The composition of the human gut microbiota is altered in many chronic human diseases. This has driven an effort to understand how microbiota contributes to disease and if it represents a target for novel therapies. In the majority of conditions, no consistent association between the disease and any specific commensal species or strain has been made, and the distinction between the microbiotas in the healthy and diseased states reflects broader differences in the community structure, diversity or density (Contijoch et al. 2019; Durack and Lynch 2019). In the rare cases where specific species or strains are hypothesized to contribute to disease, for example adhesive-invasive *Escherichia coli* (AIEC) in Crohn’s disease (Darfeuille-Michaud et al. 2004), targeted therapy using approaches such as phages may be possible (Galtier et al. 2017). In other situations, rational therapeutic approaches are elusive and the foremost strategy to modulate the microbiota has been fecal microbiota transplantation (FMT).

FMT is effective for treating recurrent *Clostridioides difficile* infection (rCDI) (van Nood et al. 2013), but is also under investigation for the treatment of ulcerative colitis (UC), Crohn’s disease (CD), liver diseases, obesity and metabolic disease, food allergy, graft-versus-host disease, check point colitis and to improve the efficacy of cancer immunotherapy (Ooijevaar et al. 2019; Wang et al. 2018; Baruch et al. 2019). Clinical trials of FMT in UC have suggested some potential for using microbiota-targeted therapy in this setting (Paramsothy et al. 2017; Moayyedi et al. 2015; Rossen et al. 2015; Costello et al. 2019), but evidence for efficacy in other indications is currently extremely limited and there is currently little or no scientific basis for the adoption of FMT as a mainstream therapeutic option outside of rCDI.

Most drugs are given with the desire of modulating a specific cellular process, aiming to restore homeostasis and health to a diseased organ or body system. As seen in rCDI and some individuals with UC, FMT can restore health to the gut but the mechanisms remain opaque. Studies in gnotobiotic mice have revealed defined gut microbiota manipulations that modulate the proportion or function of specific cells and these data are aiding the development of candidate live biotherapeutic products (LBPs) designed to specifically target certain pathways or cells, for example increasing the proportion of regulatory T cells (Cohen et al. 2019; Atarashi et al. 2013). However, to develop microbiota-targeted therapeutics for broad use it is imperative to understand the specifics of how microbiota manipulations alter host processes and phenotypes in predictable ways. Only by doing this can we hope to achieve the maximum potential benefit of these approaches while exposing treatment recipients to the minimum of risk, as we demand of all other drugs and therapeutics in common use.

This is an extremely complex problem as numerous aspects of the immune system are shaped by the gut microbiota. This is evident in gnotobiotic mice, where the gut immune system is reproducibly altered by microbiotas of different composition (Britton et al. 2019; Geva-Zatorsky et al. 2017). However, even under tightly controlled conditions in gnotobiotic mice, predictably linking microbiota composition to host phenotype is challenging. Certain components of the intestinal microbiota have defined immunomodulatory properties in the lamina propria when transferred to mice. For example, segmented filamentous bacteria (SFB) and some strains of *E. coli* induce T helper (Th)17 cells (Ivanov et al. 2009; Viladomiu et al. 2017), certain strains of *Klebsiella* can induce Th1 cells (Atarashi et al. 2017), some strains of *B. ovatus* induce high fecal IgA (Yang et al. 2019), and specific Clostridial strains can induce regulatory T (Treg) cells (Atarashi et al. 2013; Atarashi et al. 2011; Geva-Zatorsky et al. 2017). The context within which individual microbes colonize a host can alter their function. For example, immunomodulatory properties of individual strains can be altered in an inflamed host, or when co-colonized with different organisms (Lengfelder et al. 2019; Yang et al. 2019). Thus, the immune landscape of the gut is influenced by the presence of specific strains and the compositional context of the broader microbiota, making the rational design of therapies targeting the microbiota challenging.

In ex-germ free mice colonized with different complex human microbiotas from numerous human donors, population-scale associations can be made between characteristics of the microbiota donors and the phenotype transferred to mice. For example, we have previously observed that microbiotas from donors with inflammatory bowel disease induce more gut Th17 cells and fewer RORγt^+^ Treg cells than microbiotas from healthy donors (Britton et al. 2019). These altered immune populations in wild-type mice are highly predictive of disease severity in a model of colitis using ex-germ free mice colonized with the same IBD and non-IBD gut microbiotas, implying a strong functional relevance of these cells. These results suggest different compositions of gut microbiota set stable immune landscapes in the gut lamina propria and alter disease susceptibility. Specifically, the inflammatory tissue immune program in ex-germ free mice colonized by the gut microbiota of individuals with IBD and characterized by high Th17 cells and low RORγt^+^ Treg cells therefore represents a colitis-susceptible state that is potentially correctable by gut microbiota manipulation.

We established a gnotobiotic mouse experimental system to evaluate potential live bacterial therapy for IBD. This system allowed us to track recipient and donor strains during the procedure and monitor immunological changes in the host following treatment. Germ free mice were initially colonized with a defined, cultured microbiota obtained from one of three donors with IBD, followed 3 weeks later by a transplant with one of five defined consortia of healthy-donor derived microbes. In testing 14 different IBD-healthy donor combinations, we find that the inflammatory gut immune landscape induced by human IBD microbiotas is reversible and transplantation with healthy donor microbiotas can restore a tolerant immunological state. Healthy donor microbiota transplant consistently and reproducibly increased the proportion of gut RORγt^+^ Treg cells in mice initially colonized with IBD donor microbiotas. In mice colonized with an IBD donor that potently induces Th17 cells, transplants reduced the proportion of mucosal Th17 cells. We identify two distinct mechanisms associated with the recovery of gut immune health following transplant. Transplant-induced RORγt^+^ Treg cell induction was associated with increased fecal microbiota density, a measure previously associated with health (Contijoch et al. 2019; Vandeputte et al. 2017; Vieira-Silva et al. 2019). Where we observed reduced mucosal Th17 cells, this was associated with decreased colonization by a specific strain from the IBD microbiota. Further characterization revealed this strain was required for Th17 induction by this IBD microbiota. We demonstrate that the proinflammatory immune tone driven by IBD microbiotas is reversible by microbiota transplant.

### Defined microbiota transplant broadly alters gut microbiome structure

To establish a recipient-specific baseline immune tone, we first colonized germ free C57Bl/6 mice with defined, cultured fecal microbiotas (Britton et al. 2019) isolated from one of three different individuals with IBD (Figure 1A). Three weeks later, we introduced one of five defined cultured healthy donor fecal microbiota via a single gavage (i.e., a defined microbiota transplant; DMT) whereupon mice were individually housed to prevent mouse-to-mouse transmission of the DMT. We tested combinations of three IBD and five healthy donors in a minimum of six mice (14 total combinations; Figure 1A). We focused these experiments on cultured consortia where each species has a complete genome sequence, as this facilitates strain-level tracking of engraftment and downstream identification of immunomodulatory strains.

**Fig. 1.**
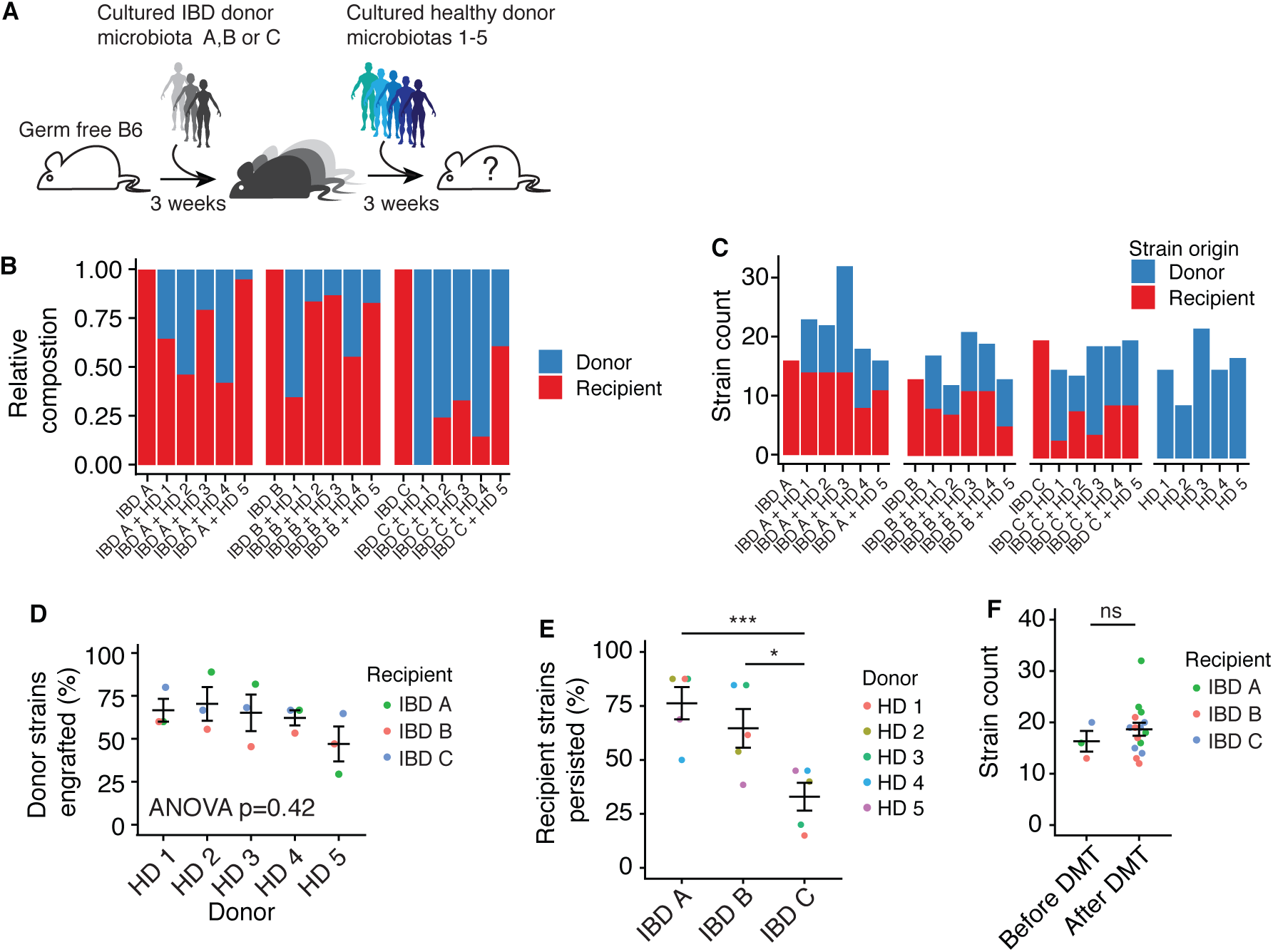
Defined microbiota transplant alters gut microbiome structure in gnotobiotic mice. (**A**) Germ free mice were first colonized with one of 3 defined IBD donor microbiotas before microbiota transplant with one of five defined healthy donor microbiotas three weeks later. (**B**) The average contribution of IBD donor and healthy donor-derived strains to the microbiota 3 weeks after transplant as a proportion of the total community. (**C**) The average number of IBD donor and healthy donor-derived strains that comprise the microbiota 3 weeks after transplant. (**D**) The average proportion of strains from each healthy donor that engraft when given as a microbiota transplant to each IBD microbiota-colonized recipient. (**E**) The average proportion of strains from each IBD donor that persist following each microbiota transplant. (**F**) The average total number of strains comprising the microbiota of mice colonized with IBD donor derived microbiota before and after transplant. Plots in D, E and F show the mean of each group of mice +/- the standard error. *p<0.05, ***p<0.001, ns – not significant by ANOVA with Tukey correction.

We performed metagenomic sequencing of DNA isolated from feces of mice before DMT, three weeks after DMT and from mice colonized with only the healthy microbiotas used in the DMTs. The healthy donor microbiotas were comprised of between 9 and 22 strains (mean 15.6). An average of 59.3% (range 31.2 – 100%) of the healthy donor strains were detected in the mice three weeks after DMT (Fig 1B and C). The proportion of strains that engrafted from each heathy donor was not significantly different (p = 0.42; ANOVA; Fig 1D) and there was no difference in the mean proportion of strains that engrafted from the donors in mice colonized with each IBD donor microbiota (p=0.18; ANOVA, Fig 1C). The three IBD microbiotas comprised 13, 17 and 20 strains respectively. An average of 58.0% (range 15.0 - 87.5%) of the strains from the IBD donors remained detectable in the mice following DMT (Fig 1E). No one donor induced a greater loss of IBD microbiota strains than any other (p = 0.98; ANOVA), but the proportion of strains from donor IBD C persisting following transplant was lower than microbiotas IBD A and IBD B (Fig 1E). On average, the number of strains lost from the microbiotas was balanced by the engraftment of healthy donor-derived strains, and the total number of strains comprising the post-transplant microbiotas was not significantly different from the number of strains detected in the IBD microbiotas before DMT (Fig 1F).

### Modulation of intestinal Th17 cells by DMT

Three weeks after DMT, we assessed the impact of the transplanted microbes on the host intestine tissue resident immune populations in comparison to mice receiving no transplant that were colonized with either recipient IBD microbiota or healthy donor microbiota alone. We used flow cytometry to profile CD4 T cell populations in the colon and ileum for each donor, recipient, and donor+recipient combination. A hallmark of mice colonized with IBD donor microbiotas is an increase in the proportion of gut Th17 cells, relative to mice colonized with healthy donor microbiotas (Britton et al. 2019; Viladomiu et al. 2017; Atarashi et al. 2015). As demonstrated in multiple animal models, increased Th17 cells are associated with susceptibility to inflammatory disease (Miossec and Kolls 2012). Therefore, we hypothesize that a rational aim of microbiota-targeted therapy for IBD is to develop interventions that reduce these potentially pathogenic microbiota-induced Th17 cells. With this in mind, we selected as DMT donors healthy donor-derived microbiotas that consistently induced a low proportion of LP Th17 cells (Fig 2, A). We measured the proportion of gut Th17 cells in mice colonized with IBD microbiotas alone, healthy donor mice alone and in mice with IBD microbiotas that had received healthy DMT. IBD microbiota A, from a donor with Crohn’s disease, was notable for inducing a particularly high proportion of colonic Th17 cells in gnotobiotic mice (Fig 2, B) (Britton et al. 2019). In mice colonized with microbiota IBD A, DMT with all of the healthy microbiotas reduced the proportion of Th17 cells in the colon (Fig 2, B and C). In mice colonized with IBD B or IBD C where baseline Th17 cells was lower, DMT did not significantly reduce the proportion of colon Th17 cells. A similar effect was observed in the ileum where the proportion of Th17 cells was reduced following four of the five DMTs in mice colonized with IBD A, but not significantly altered in mice colonized with IBD B or C (Fig. S2).

**Fig. 2.**
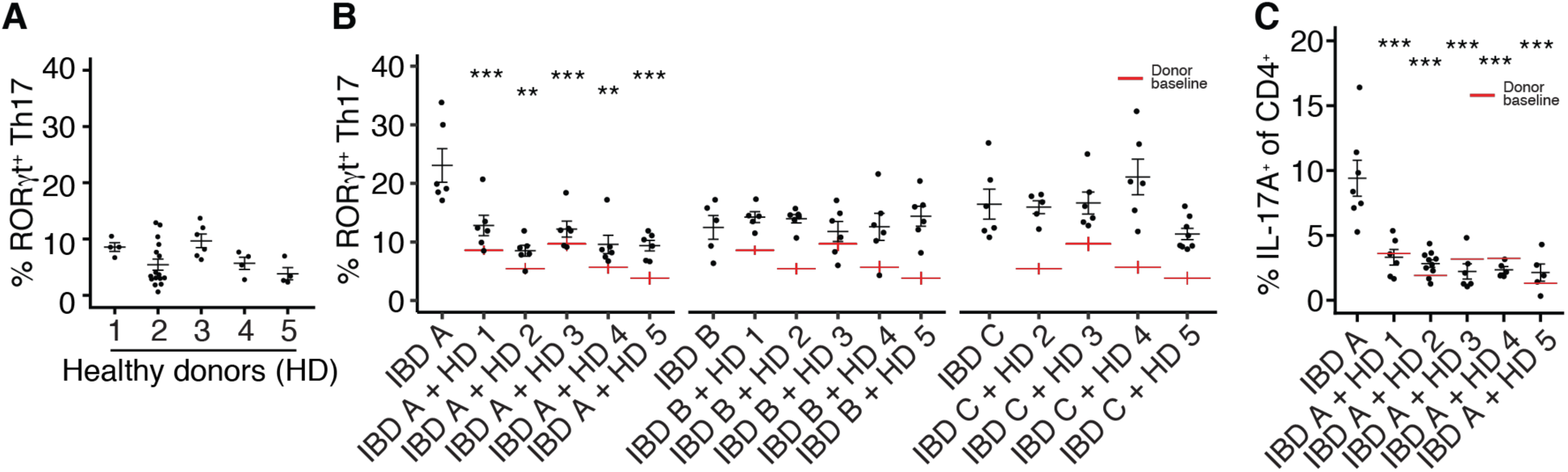
Modulation of mucosal Th17 cells by defined microbiota transplant. **(A)** The proportion of RORγt^+^ Th17 cells (of live, CD4^+^FoxP3^-^cells) in the colon lamina propria of gnotobiotic mice colonized with the five HD-derived microbiotas used as DMT donors. (**B**) The proportion of colon lamina propria RORγt^+^ Th17 cells (of live, CD4^+^FoxP3^-^cells) in groups of mice colonized with each IBD donor alone or three weeks following DMT with one of the five HD microbiotas. Red lines indicate the proportion of Th17 cells induced by each HD alone, as shown in (A). (**C**) The proportion of IL-17A^+^ CD4^+^ T cells (of live, CD4^+^ cells) in the colon lamina propria of mice colonized with IBD A alone, or three weeks following DMT with one of the five HD microbiotas. Each point represents data from one mouse and black lines indicate the mean and standard error of each group. Red lines indicate the mean proportion of the specified cells induced by each HD alone. **p<0.01, ***p<0.001, ****p<0.0001 as assessed by ANOVA with Tukey correction.

### Identification of a Th17-inducing strain from a donor with Crohn’s disease

Intestinal Th17 cells can be induced by specific microbes, for example segmented filamentous bacteria (SFB) (Ivanov et al. 2009), a strain of *Bifidobacterium adolescentis* (Tan et al. 2016) and specific strains of *Escherichia coli* (Viladomiu et al. 2017). As the proportion of Th17 cells induced by donor IBD A was particularly high, we hypothesized that the induction of Th17 cells by this microbiota was due to the presence of a specific strain that drives their induction and that perhaps the abundance of this strain was modulated following DMT, leading to the reduction in Th17 cells.

Using 16 strains from the IBD A microbiota, we assembled eight sub-communities, each of four strains, using an orthogonal design (Faith et al. 2014). Each strain was present in two different sub-communities. We colonized groups of germ-free B6 mice with the eight sub-communities and assessed the impact of each on the proportion of colon Th17 cells (Fig 3, A). The eight sub-communities induced varied proportions of IL-17A^+^ Th17 cells (p=0.0004, ANOVA; Fig 3, B). We quantified the association of each strain with the proportion of Th17 cells and found a single strain, Escherichia coli ‘A6’, positively associated with the proportion of colon RORγt^+^ and IL-17A^+^ Th17 cells (p=0.0035, t-test; Fig 3, C and Fig. S3). Of note, a second strain of *E. coli*, isolated from IBD A (‘E2’) was not associated with Th17 cell in this screen (p=0.94, t-test, Fig S3). To confirm the specific ability of *E. coli* A6 to induce Th17 cells we generated a consortium of the strains isolated from IBD donor A excluding only *E. coli* A6. Mice colonized with this consortium lacking only *E. coli* A6 (‘ΔA6’) had a significantly reduced proportion of colon IL-17A^+^ Th17 cells compared to mice colonized with the complete community from donor IBD A (Fig 3, D). CD4^+^ T cells from the mLN of mice colonized with IBD A secreted IL-17A when restimulated *ex vivo* with dendritic cells loaded with a lysate of *E. coli* A6 (Fig 3, E) (Yang et al. 2014). This response was prevented by MHC-II blockade, suggesting a cognate interaction between the Th17 cells and DC presenting antigen from *E. coli* A6 (Fig 3, E). These data demonstrate that strain *E. coli* ‘A6’ is a dominant inducer of antigen-specific Th17 cells in the microbiota of donor IBD A.

**Fig. 3.**
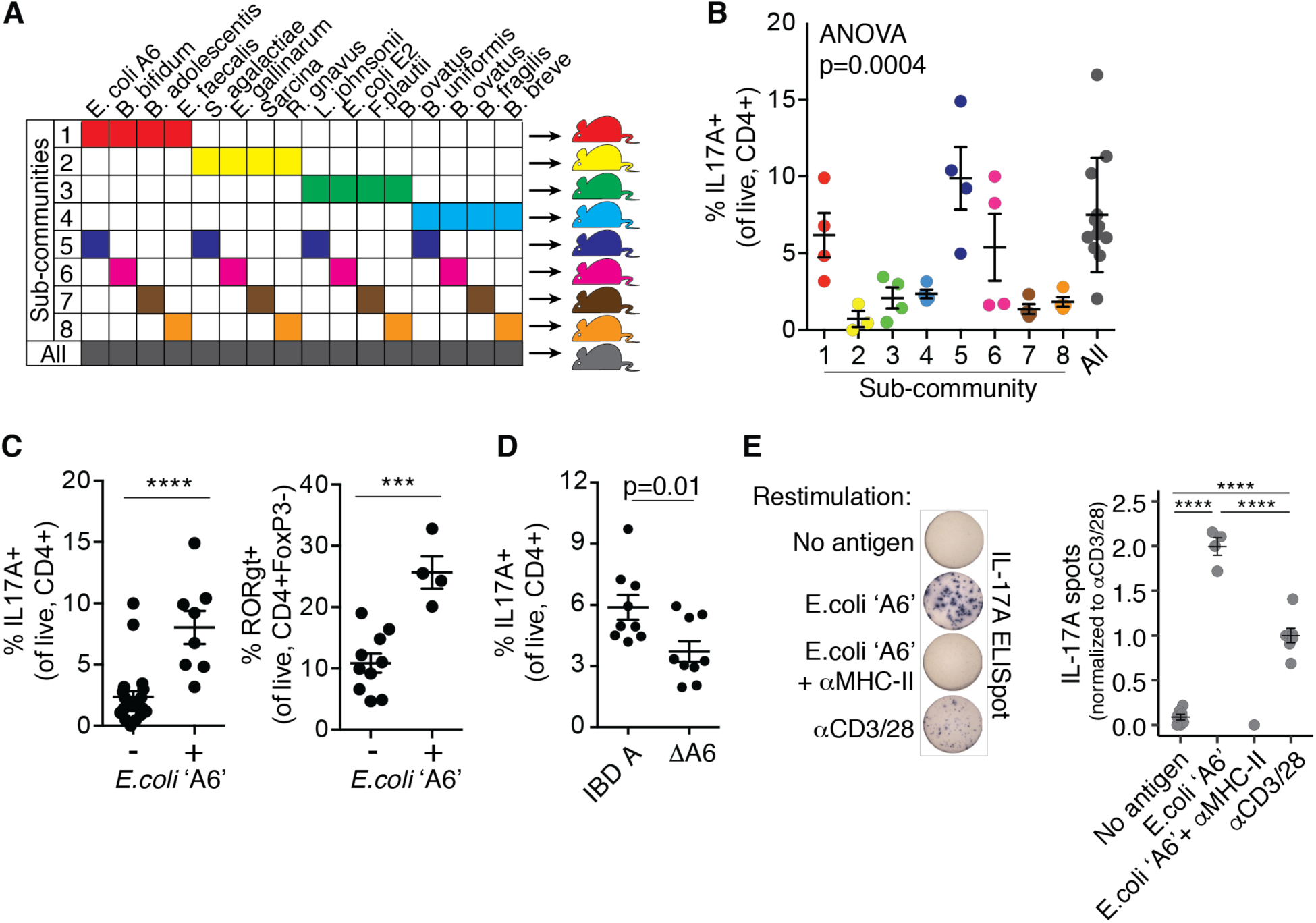
Identification of a Th17-inducing strain from a donor with Crohn’s disease. **(A)** Groups of germ free B6 mice were each colonized with one of eight communities comprised of 4 of the 16 strains isolated from donor IBD A (**B**) The proportion of IL-17A^+^ CD4^+^ T cells (of live, CD4^+^ cells) in the colon lamina propria of mice colonized with each sub-community derived from the IBD A microbiota. (**C**) The proportion of IL-17A^+^ CD4^+^ T cells and RORγt^+^ Th17 cells in mice colonized with communities in which E. coli strain A6 was present (+) or absent (-). (**D**) The proportion of IL-17A^+^ CD4^+^ T cells (of live, CD4^+^ cells) in the colon lamina propria of mice colonized with the complete IBD A microbiota or a modified version of IBD A lacking E. coli strain A6 (ΔA6). (**E**) IL-17A secretion from mLN CD4^+^ T cells isolated from mice colonized with IBD A and restimulated *ex vivo* with anti-CD3 and anti-CD28 or dendritic cells loaded with E. coli strain A6, with or without an MHC-II blocking antibody. Plots show the mean and standard error of each group of mice and P values were calculated by t-test (C and D), ANOVA (B) and ANOVA with Tukey correction (E). ***p<0.001, ****p<0.0001

We analyzed the relative abundance of the IBD-donor derived strains in mice both before and following DMT with each of the five healthy donor microbiotas. The abundance of the Th17-inducing *E. coli* ‘A6’ strain was significantly reduced in feces of mice following DMT relative to mice colonized with IBD A alone (Fig 4, A). The relative abundance of *E. coli* ‘A6’ in the fecal microbiota before and after DMT was associated with the proportion of colon RORγt^+^ Th17 cells (Fig 4, B). This suggests a mechanism whereby, in the context of the IBD A microbiota, DMT with healthy donor microbiotas reduces the relative abundance of a specific strain, resulting in a reduction in lamina propria Th17 cells.

**Fig. 4.**
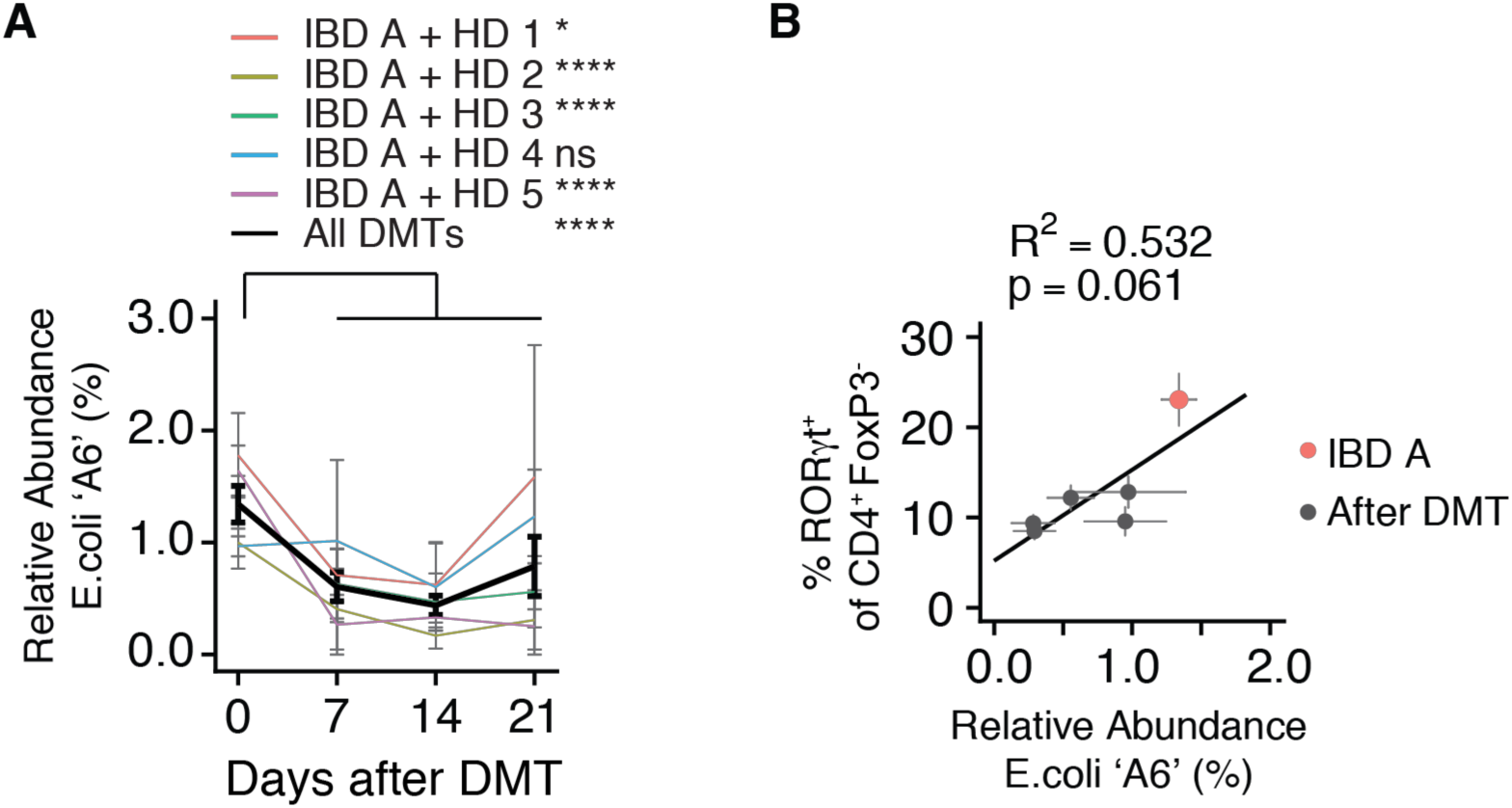
Modulation of an IBD-associated Th17-inducing strain following microbiota transplant. **(A)** The mean relative abundance of E coli strain A6 from donor IBD A in mice before and at three timepoints after DMT with one of five HD microbiotas. (**B**) The correlation between the relative abundance of E. coli strain A6 and the proportion of RORγt^+^ Th17 cells in mice colonized with IBD A alone, or following transplant with each of the five HD microbiotas. *p<0.05, ****p<0.0001 as calculated by ANOVA with Tukey correction comparing the relative abundance of the strain before transplant and at each timepoint after transplant. P value in (B) calculated by F-test.

### Induction of regulatory T cells following DMT

The RORγt^+^ subset of intestinal regulatory T cells enforces tolerance to microbiota (Ohnmacht et al. 2015). RORγt^+^ Treg cells are dynamically and variably induced by different microbiotas (Sefik et al. 2015) and are specifically deficient in mice colonized with microbiotas from donors with IBD (Britton et al. 2019). We hypothesize that one rational aim of microbiota-targeted therapy for IBD would be to increase the *in situ* differentiation and stability of RORγt^+^ Treg cells. When colonized alone, each of the three IBD recipient microbiotas (IBD A-C) induced a relatively low proportion of RORγt^+^ Treg cells in both colon and ileum of mice (Fig 5, A and B). Three weeks following microbiota transplant, the proportion of RORγt^+^ Treg cells in the colon of recipient mice was in all cases significantly increased relative to mice colonized with each IBD microbiota alone (Fig 5, A). We observed no “super donor” microbiota that consistently induced a higher proportion of RORγt^+^ Treg cells across all recipient microbiotas, and mice colonized with each of the three IBD donor microbiotas were similarly amenable to immune modulation by DMT in the colon.

**Fig 5.**
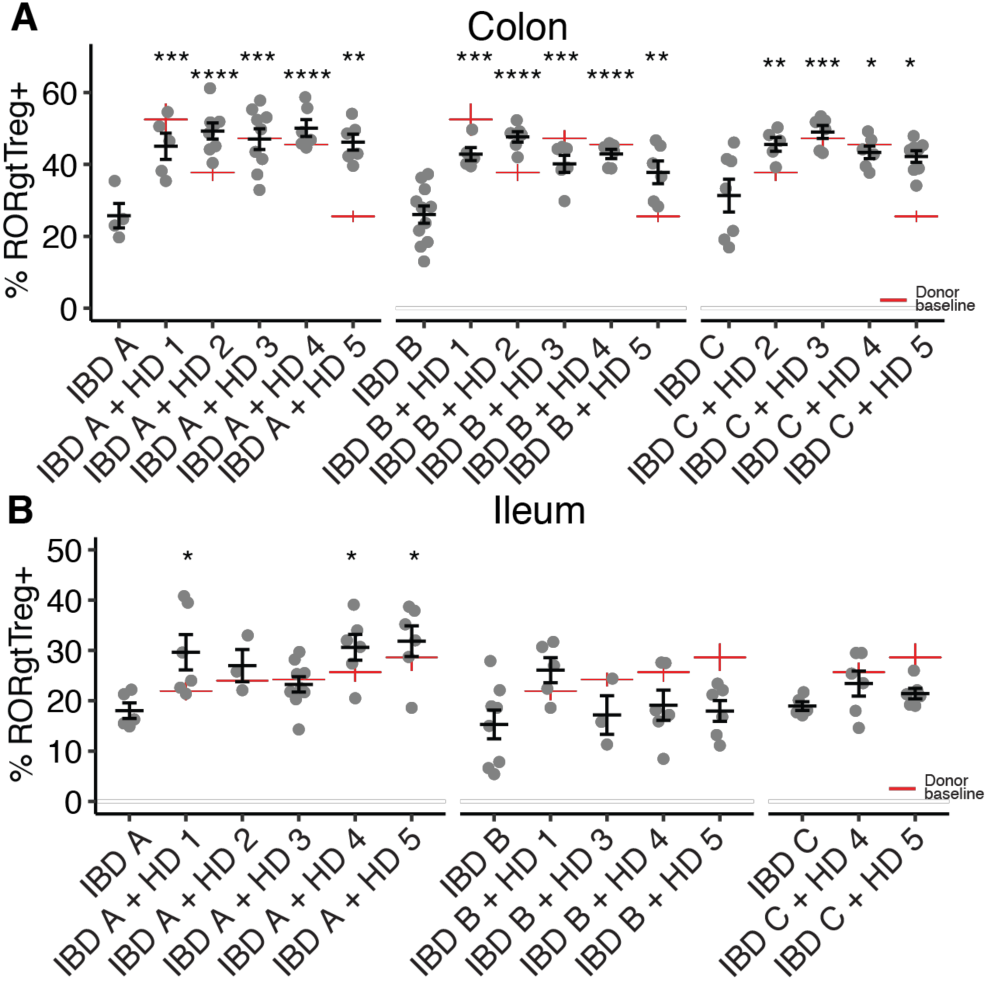
RORγt^+^regulatory T cells are induced following defined microbiota transplant. **(A-B)** The proportion of colon and ileum lamina propria RORγt^+^ Treg cells (of live, CD4^+^FoxP3^+^ cells) in groups of mice colonized with each IBD donor alone or three weeks following DMT with one of the five HD microbiotas. Red lines indicate the proportion of RORγt^+^ Treg cells induced by each HD alone. Plots show the mean and standard error of each group of mice. *p<0.05, **p<0.01, ***p<0.001, ****p<0.0001 by ANOVA with Tukey correction.

Modulation of RORγt^+^ Treg cells was less pronounced in the ileum where significant increases were limited to three of the DMTs only in mice colonized with IBD A (Fig 5, B). In addition to the RORγt^+^ subset of Treg cells, we also observed modulation of the total gut FoxP3^+^ Treg population (Fig S5, A and B). Microbiotas that induce high proportions of RORγt^+^ Treg cells are also associated with reduced activation of colon dendritic cells (Britton et al. 2019; Ohnmacht et al. 2015). Consistent with these observations, the expression of CD86 was lower on both total and CD11b^+^CD103^+^ (double positive) colonic dendritic cells from mice colonized with HD 1 than IBD A and are reduced post-DMT (Fig S5, C).

The proportion of total FoxP3^+^ Treg in colon and ileum was less consistently altered following DMT (Fig S5, A,B). Interestingly, the proportion of colon FoxP3^+^ Treg cells in each recipient+donor microbiota combination was predictable from the baseline induction by the donor microbiota alone (R^2^ = 0.47; p = 0.001, f-test) (Fig S5, D). However, the proportion of RORγt^+^ Treg cells was not predictable from the baseline induction of these cells by the donor and recipient (Fig S5, E). The observations of RORγt^+^ Treg cell induction following every tested DMT, and that this was unrelated to the proportion induced by the donors alone suggested that RORγt^+^ Treg cell induction was occurring by a mechanism unrelated to the specific composition of the microbiotas and may relate to broader structural characteristics of the communities.

### Microbiota density increases following microbiota transplant and is associated with increased mucosal Treg cells in mice and endoscopic remission in individuals with UC

The IBD microbiota has reduced alpha diversity (Morgan et al. 2012; Manichanh et al. 2006) and lower density than the microbiota from healthy donors (Contijoch et al. 2019; Vandeputte et al. 2017; Frank et al. 2007; Vieira-Silva et al. 2019). The proportion of intestinal Treg cells can be influenced by microbiota composition (Britton et al. 2019; Atarashi et al. 2013; Atarashi et al. 2011; Geva-Zatorsky et al. 2017; Sefik et al. 2015), but can also be regulated independent of composition in response to changes in microbiota density (Contijoch et al. 2019). Following FMT for rCDI in humans, both the alpha diversity and the density of the gut microbiota is increased (Contijoch et al. 2019; Song et al. 2013; Khanna et al. 2017; Shankar et al. 2014; Kellingray et al. 2018). We therefore examined if the increase in RORγt^+^Treg cells we observe following DMT in our model could be attributed to changes in either alpha diversity or microbiota density (Fig 6). We calculated microbiota density as the amount of DNA extracted from a fecal sample divided by the original sample mass (Contijoch et al. 2019; Reyes et al. 2013). Alpha diversity increased following some of the DMTs, but this was not consistent for all tested pairs of IBD and healthy donor microbiotas (p=0.07, paired t test; Fig 6, A and B). However, microbiota density was significantly increased after DMT (p=0.004, paired t-test; Fig 6,C and D.

**Fig. 6.**
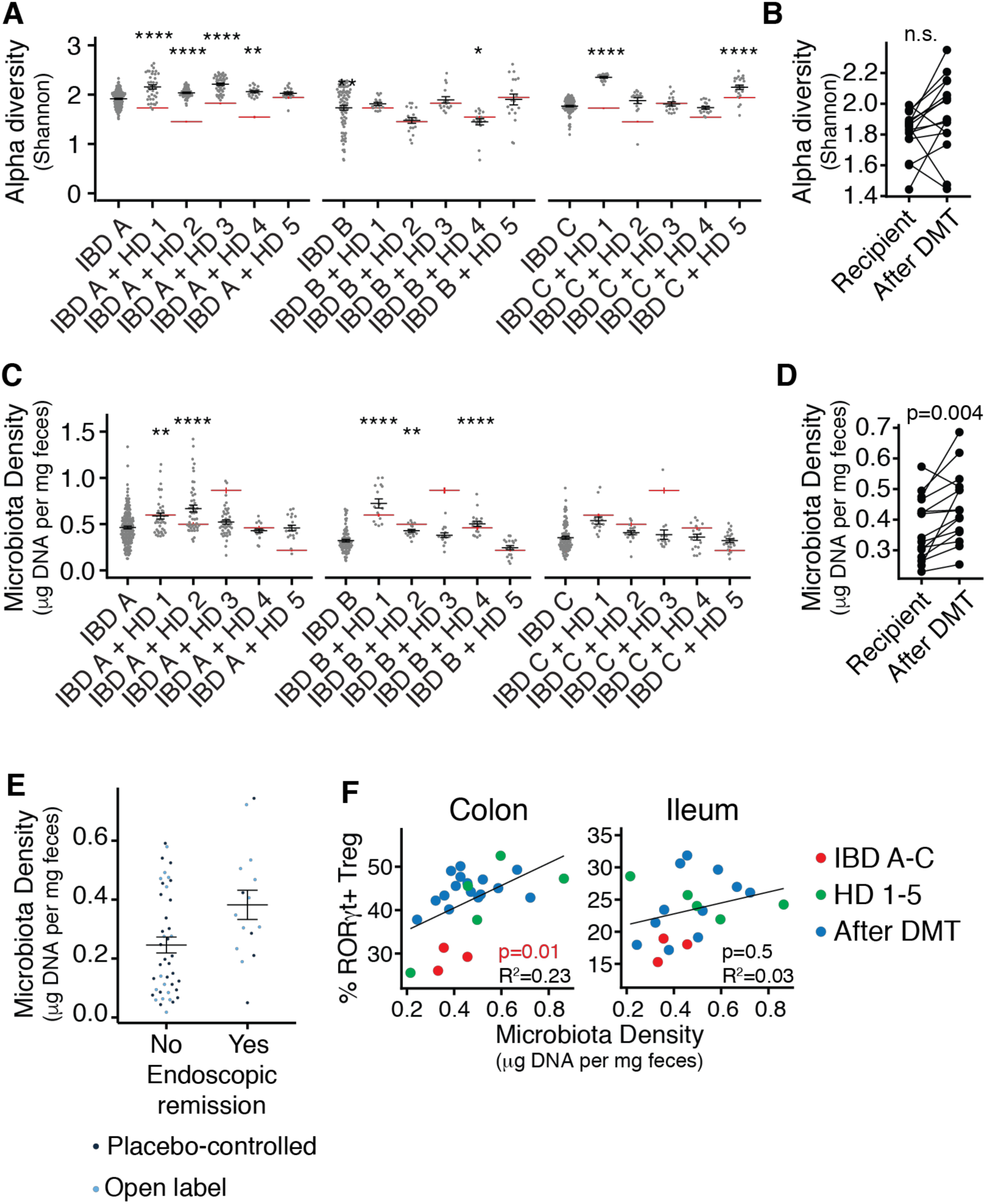
Microbiota density is increased following defined microbiota transplant and is associated with increased mucosal Treg cells in mice and endoscopic remission in individuals with UC following FMT. **(A)** The mean difference in fecal microbiota density in mice colonized with IBD microbiotas before and after DMT. (**B**) Fecal microbiota density in groups of mice colonized with each IBD donor alone or three weeks following DMT with one of the five HD microbiotas. Red lines indicate the fecal microbiota density in mice colonized with HD microbiotas alone. **(C)** Alpha diversity of fecal microbiota in groups of mice colonized with each IBD donor alone or three weeks following DMT with one of the five HD microbiotas. Red lines indicate the alpha diversity of fecal microbiota in mice colonized with HD microbiotas alone. (**D**) The mean difference in alpha diversity in mice colonized with IBD microbiotas before and after DMT. (**E**) Fecal microbiota density of FMT clinical trial participants with ulcerative colitis who did and did not achieve endoscopic remission following treatment whether in the placebo controlled or open label arm of the trial. (**F**) The association between fecal microbiota density and the proportion of mucosal RORγt^+^ Treg cells in mice colonized with HD and IBD donor microbiotas alone or following DMT. in A and C *p<0.05, **p<0.01, ****p<0.0001 by ANOVA with Tukey correction. P values in B and D calculated by paired t-test and in F by f-test.

Increased fecal microbiota density has been reported in individuals receiving FMT to treat rCDI (Contijoch et al. 2019). To assess the relevance of microbiota density modulation in FMT used in the context of IBD, we analyzed data from a recent placebo-controlled clinical trial of FMT in individuals with ulcerative colitis (Paramsothy et al. 2017). This study reported endoscopic remission in ∼20% of individuals receiving an intensive multi-donor FMT regime. Strikingly, microbiota density was higher 8 weeks following an 8-week intensive FMT therapy in individuals in which endoscopic remission was achieved than in those where FMT did not lead to endoscopic remission (p = 0.024; Welch two sample t-test, Fig 6, E).

In the gnotobiotic mouse model, microbiota density but not alpha diversity was significantly associated with the proportion of colonic RORγt^+^ Treg and total FoxP3^+^ Treg cells (R^2^ =0.23, p=0.01 and R^2^ =0.14, p=0.01 respectively, f test; Fig 6, F and Fig 6S, A and B). A similar effect was observed in mice colonized with an SPF microbiota and treated with different antibiotics to modulate microbiota density. The greater the depletion of microbiota density following antibiotic treatment the greater the reduction of mucosal RORγt^+^ Treg (R^2^ = 0.49, p<0.0001, R^2^ = 0.11, p=0.02 for colon and ileum respectively; Fig S6, D). Taken together, these data suggest that DMT with healthy donor microbiotas can increase microbiota density in mice colonized with IBD donor microbiotas and in some humans receiving FMT for UC. In mice, the restoration of microbiota density correlates with the induction of gut Treg populations and in humans with UC it is associated with endoscopic remission, suggesting reinstatement of tolerance in the intestinal tissues following transplantation.

## Discussion

Compositional changes to the gut microbiota are observed in many human diseases. Fecal microbiota transplantation is predicated on the assumption that an altered microbiota contributes to disease and restoring the composition of the microbiota to a “normal” state will provide therapeutic benefit. Despite a steady increase in FMT trials for different indications, we have little data to make predictions about how a treatment recipient will respond to a microbiota transplant, what are desirable traits of a good donor, and molecular markers to indicate success beyond standard clinical endpoints. Manipulation of microbiota community characteristics, including colonization resistance, microbiota fitness, and microbiota density are likely key to the success of FMT in the context of recurrent *C. difficile* infection. However, the application of FMT in the context of autoimmune or inflammatory diseases likely requires achieving an element of specific immune modulation either in the gut, at a distal tissue site, or systemically. Immune modulation directed by microbiota manipulation could be achieved through a combination of suppression or elimination of proinflammatory effector strains and expansion or addition of tolerance-promoting strains or community characteristics. To advance microbiota manipulation as a potential therapeutic for immune mediated disease, and to provide a solid scientific basis for the expanded use of such therapies beyond rCDI, we must better understand the potential of microbiota manipulation to alter specific immune populations and ultimately track these immune populations in FMT clinical trials (Vaughn et al. 2016; Zhang et al. 2016; Jacob et al. 2017).

We report a set of gnotobiotic mouse experiments that support the pre-clinical assessment of LBPs and microbiota transplants in the context of human inflammatory disease-associated microbiotas. In this study, we have used mice colonized with defined IBD donor microbiotas, and assessed how the immunophenotype of these hosts responds to DMT with healthy donor microbiotas with known immunomodulatory properties across a total of 14 different microbiota transplants. The gut immune landscape of the mice is consistently reshaped following DMT. We demonstrate that introduction of healthy donor microbiotas to mice colonized with IBD microbiotas can lead to an increase in the proportion of Treg cells, including the potently immunoregulatory RORγt^+^ Treg subset. We find that the increase in RORγt^+^ Treg cells occurs with a restoration of microbiota density following DMT. Microbiota density is decreased in humans with IBD, and is correlated with the proportion of gut Treg cells in specific pathogen free mice (Contijoch et al. 2019; Vandeputte et al. 2017). We find increased microbiota density in individuals with ulcerative colitis reaching endoscopic remission following intensive FMT in a recent placebo-controlled clinical trial (Paramsothy et al. 2017), relative to individuals where endoscopic remission did not occur. We therefore hypothesize that the DMTs in mice are restoring the microbiota to a compositional or structural state that supports a greater microbiota density and this leads to induction of RORγt^+^ Treg cells. In humans receiving FMT for ulcerative colitis, a higher microbiota density following treatment associates with better endoscopic outcomes, suggesting a shift towards a more tolerance-promoting mucosal environment. Interestingly, the increase in RORγt^+^ Tregs was not correlated with increases in alpha diversity, which is frequently associated with health and is one of the most commonly used metrics of ‘dysbiosis’. This suggests that microbiota density may be a metric of microbiota health as it relates to the tolerogenic Treg compartment, and could be a useful biomarker for non-invasive monitoring of fecal microbiota transplant (Contijoch et al. 2019).

Microbiotas from donors with IBD induce a greater proportion of Th17 cells in the gut than healthy donor microbiotas (Britton et al. 2019; Viladomiu et al. 2017). Focusing on mice colonized with a microbiota from a donor with Crohn’s disease, we found that DMT using healthy donor microbiotas suppressed the proportion of gut Th17 cells. This reduction in Th17 cells following DMT was not consistently associated with changes in microbiota density or alpha diversity. Instead, we found that a specific strain of *E. coli* was a primary driver of Th17 cell enrichment in mice colonized with this donor consortia and decreased Th17 cells following DMT was correlated with a significant decrease in the abundance of this strain. Whereas RORγt^+^ Treg cell proportions were modulated by DMT following broad changes to the microbiota density, the reduction of Th17 cells following MT in mice colonized with this CD donor microbiota may be attributed to depletion of one specific strain. This more broadly reflects different rationale being used to design microbiota-based therapeutics. Some approaches, such as FMT, aim to reverse ‘dysbiosis’ and restore a broad structural and compositional state that is associated with health. Other approaches, such as the use of phage therapy or CRISPR/Cas-based strategies (Cohen et al. 2019; Galtier et al. 2017) aim to target specific strains implicated in the particular disease. If microbiota-targeted therapies are to be successfully used in the context of complex diseases beyond rCDI it is likely that a combination of these strategies will be required.

## Methods

### Human microbiota consortia

Microbiota HD 5 is a previously described consortia of predominantly Clostridial species (Atarashi et al. 2013; Narushima et al. 2014). Cultured consortia of microbes from human donor fecal samples IBD A-C and HD 1-4 were prepared as previous described (Britton et al. 2019). Briefly, a variety of solid media were inoculated with a slurry of each stool sample and following incubation under anaerobic and aerobic conditions, liquid media (LYBHIv4 (Sokol et al. 2008); 37 g/l Brain Heart Infusion [BD], 5g/l yeast extract [BD], 1 g/l each of D-xylose, D-fructose, D-glactose, cellubiose, maltose, sucrose, 0.5 g/l N-acetylglucosamine, 0.5 g/l L-arabinose, 0.5 g/l L-cysteine, 1g/l malic acid, 2 g/l sodium sulfate, 0.05% Tween 80, 20 μg/ml menadione, 5 mg/l hemin (as histidine-hemitin), 0.1 M MOPS, pH 7.2) arrayed in multiwell plates was inoculated with the resulting colonies and cultures maintained under anaerobic conditions. Strains comprising each consortia were characterized by a combination of MALDI-TOF mass spectrometry, 16S rDNA amplicon and whole genome shotgun sequencing. Strains were pooled in equal volumes and stored at −80°C with 15% glycerol and defrosted immediately before inoculating mice.

### Mice and gnotobiotic methods

Germ free C57Bl/6J mice were bred in isolators at the Mount Sinai Precision Immunology Institute Gnotobiotic Facility. SPF C57Bl/6J mice were purchased from Jackson Laboratories. Mice were colonized at approximately six weeks old by a single gavage of a defined microbiota consortia. Following colonization, mice were housed in autoclaved filter-top cages, fed autoclaved diet (5K67, LabDiet) and water, and handled under aseptic conditions. Some mice received a defined microbiota transplant, administered as a single gavage three weeks after initial primary colonization. At the time of DMT, mice were transferred to a new autoclaved cage to minimize the carryover of the pre-existing microbiota from the cage environment. Where indicated, mice were provided antibiotics at the following concentrations; ampicillin 1 mg/ml, ciprofloxacin 0.1 mg/ml, clindamycin 0.267 mg/ml, polymyxin B 0.1 mg/ml or vancomycin 0.5 mg/ml in drinking water with 2% sucrose, prepared as previously described (Contijoch et al.2019).

### Lymphocyte isolation

Gut tissues were prepared are previously described (Britton et al. 2019). Briefly, cleaned gut tissues were deepithelialized in 5mM EDTA, 15mM HEPES and 5% FBS in HBSS before digestion with 0.5mg/ml Collagenase Type IV (Sigma Aldrich) and 0.25mg/ml DNase 1 in HBSS with 2% FBS. Lymphocytes enriched by passing the cell suspension sequentially though 100 μm and 40 μm strainers. For EliSPOT experiments, CD4^+^ T cells were isolated from mLN using magnetic isolation (CD4 microbeads, Miltentyi Biotech) and dendritic cells were isolated from the spleens of naïve SPF C57B/6 mice (Jackson Laboratories) by magnetic isolation (CD11c microbeads, Miltenyi Biotech) using an AutoMACS instrument (Miltenyi Biotech) following the manufacturer’s instructions.

### Flow cytometry

Flow cytometry of CD4^+^ T cells was performed as previously described (Britton et al. 2019). The following antibodies were used for T cell phenotyping: CD45 APC-Cy7 (Biolegend), CD4 APC (Biolegend), FoxP3 PE (Thermo Fisher/eBioscience), RORgt PerCp-Cy5.5 (BD Bioscience), IL-17A-PE (Biolegend) and dead cells were excluded using Zombie Aqua (Biolegend). To detect intracellular IL-17A, isolated lymphocytes were first restimulated with 5 ng/ml phorbal 12-myristate 13-acetate (PMA) and 500 ng/ml ionomycin in the presence of monensin (Biolegend) for 3.5 hours. FoxP3 and RORγt expression was analyzed in unstimulated cells. For analysis of dendritic cells, the following antibodies were used: CD45 BrilliantViolet 750 (BD Bioscience), MHC-II Pacific Blue (Biolegend), CD11b PerCP-Cy5.5 (Biolegend), CD11c PE-Cy7 (Biolegend), CD64 BrilliantViolet 786 (BD Bioscience), CD86 BrilliantViolet 605 (Biolegend) and dead cells were excluded with Fixable Viability Dye eFluor780 (ThermoFisher/eBioscience). T cell data was generated using an LSRII instrument (BD Biosciences) and dendritic cell phenotype data was acquired using an Aurora spectral cytometer (Cytek). Baseline immune profiles for the three recipient communities and four of the five donor communities (except Donor 5) were previously published (Britton et al. 2019) and are included here as baseline reference points to compare with the results of the DMT.

### EliSPOT Assay

EliSPOT experiments for the detection of microbe-specific T cell responses were performed as previously described with some minor modifications (Yang et al. 2014). CD4^+^ T cells were isolated from the mLN of gnotobiotic mice using magnetic isolation (CD4 microbeads, Miltentyi Biotech) and dendritic cells were isolated from the spleens of naïve SPF C57B/6 mice (Jackson Laboratories) by magnetic isolation (CD11c microbeads, Miltenyi Biotech) using an AutoMACS instrument (Miltenyi Biotech) following the manufacturer’s instructions. Bacterial strains were grown to stationary phase in LYBHIv4 media, and bacterial antigen was prepared by autoclaving the washed bacteria in PBS. Dendritic cells were plated at a density of 50,000 cells/well and pulsed with antigen (1:100 dilution of the stationary phase culture) overnight. Isolated CD4^+^ T cells were added to the culture the next day (2:1 T cell to dendritic cell ratio) and placed at 37 degrees for 48 hours. Where indicated, MHC class II blocking antibody (M5/114.15.2, Biolegend, 2.5μg/ml) was added to DC cultures one hour prior to adding the CD4^+^ T cells. After 48 hours of co-cultured, the cells were transferred to an IL-17A ELISPOT plate for an additional 24hours and then developed according to the manufacturer’s instructions (R&D Systems).

### Genome and metagenome sequencing

DNA was extracted from isolated bacterial strains by mechanical dissociation by bead beating in 282 μl of DNA buffer A (20 mM Tris pH 8.0, 2 mM EDTA and 200 mM NaCl), 200 μl of 20% SDS (v/w), 550 μl of Phenol:Chloroform:IAA (25:24:1), 268 μl of Buffer PM (Qiagen), and 400 μl of 0.1 mm diameter zirconia/silica beads (Yang et al. 2019). DNA was extracted from mouse feces by bead beating in DNase Inactivation Buffer (DIB; 0.5% SDS, 0.5 mM EDTA, 20 mM Tris (pH 8.0) with 200μl of 0.1 mm diameter zirconia/silica beads (Contijoch et al. 2019). DNA from bacterial isolates and feces was isolated using QiaQuick (Qiagen) columns and quantified by Qubit assay (Life Technologies). Sequencing libraries were generated from sonicated DNA with the NEBNext Ultra II DNA Library Prep kit (New England BioLabs). Ligation products were purified with SPRIselect beads (Beckman Coulter) and enrichment PCR performed with NEBNext Ultra Q5 Master Mix (New England BioLabs). Samples were pooled in equal proportions and size-selected using 0.6x followed by 0.2x of AMPure XP beads (Beckman Coulter) before sequencing with an Illumina HiSeq (paired-end 150 bp). Reads from metagenomic samples were trimmed, subsampled to 100,000 reads and mapped to the unique regions of bacterial genomes known to potentially form part of the microbiota in a given gnotobiotic sample. Abundances were scaled to the size of each specific genome as previous described (McNulty et al. 2013). Microbiota density was calculated as the mass of DNA extracted from a fecal sample, divided by the mass of the sample (Contijoch et al. 2019).

## Acknowledgments

We thank C. Fermin, E. Vazquez and G. N. Escano of the Mount Sinai Immunology Institute Gnotobiotic Facility and E. Ariztia, O. Vennaro and Z. Li for technical support. This work was supported in part by the staff and resources of Scientific Computing and of the Flow Cytometry Core at the Icahn School of Medicine at Mount Sinai. Sequencing data has been deposited to NCBI; Bioprojects PRJNA518912 and PRJNA589044.

## Author contributions

G.J.B, E.J.C and J.J.F. designed experiments. G.J.B, E.J.C, M.P.S., G.B. and A.B. performed experiments. G.J.B, E.J.C, V.A. and J.J.F. analysed data. L.S.M., A.D., D.G., T.J.B., N.O,K., M.A.K, H.M., S.P., J.C.C., J-F, C., M.C.D. and A.G. samples, analyses or clinical data. J.J.F. superisized the project. G.J.B. wrote the paper with input from all authors.

