## Supplemental Figures for "Defined microbiota transplant restores Th17/RORγt^+^ regulatory T cell balance in mice colonized with inflammatory bowel disease microbiotas"

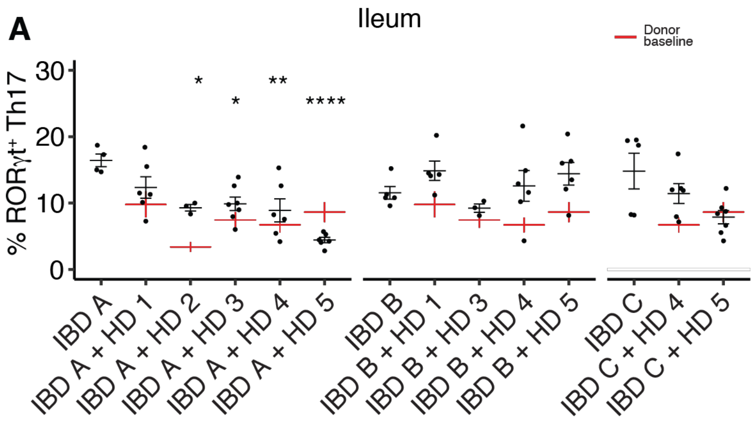


**Fig S2. Modulation of mucosal Th17 cells by defined microbiota transplant. (A)** The proportion of RORγt$^+$ Th17 cells (of live, CD4$^+$FoxP3$^-$ cells) in the ileum lamina propria of gnotobiotic mice colonized with the five HD-derived microbiotas used as DMT donors. Red lines indicate the proportion of Th17 cells induced by each HD alone. Plots show the mean and standard error of each indicated group of mice. *$p<0.05$, **$p<0.01$, ****$p<0.0001$ by ANOVA with Tukey correction.

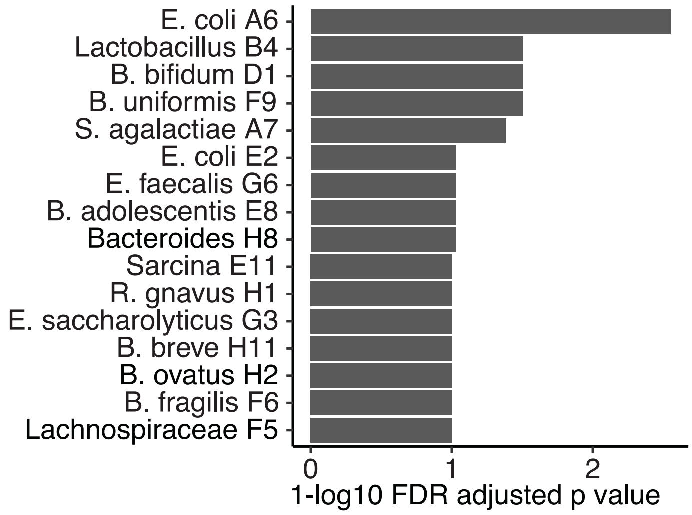


**Fig S3. Identification of a Th17-inducing strain from a donor with Crohn's disease.** FDR-adjusted one-tailed p values for the positive association of each strain from defined microbiota IBD A with the proportion of colon IL-17A$^{+}$ CD4 T cells derived from the in vivo gnotobiotic screen described in Figure 3.

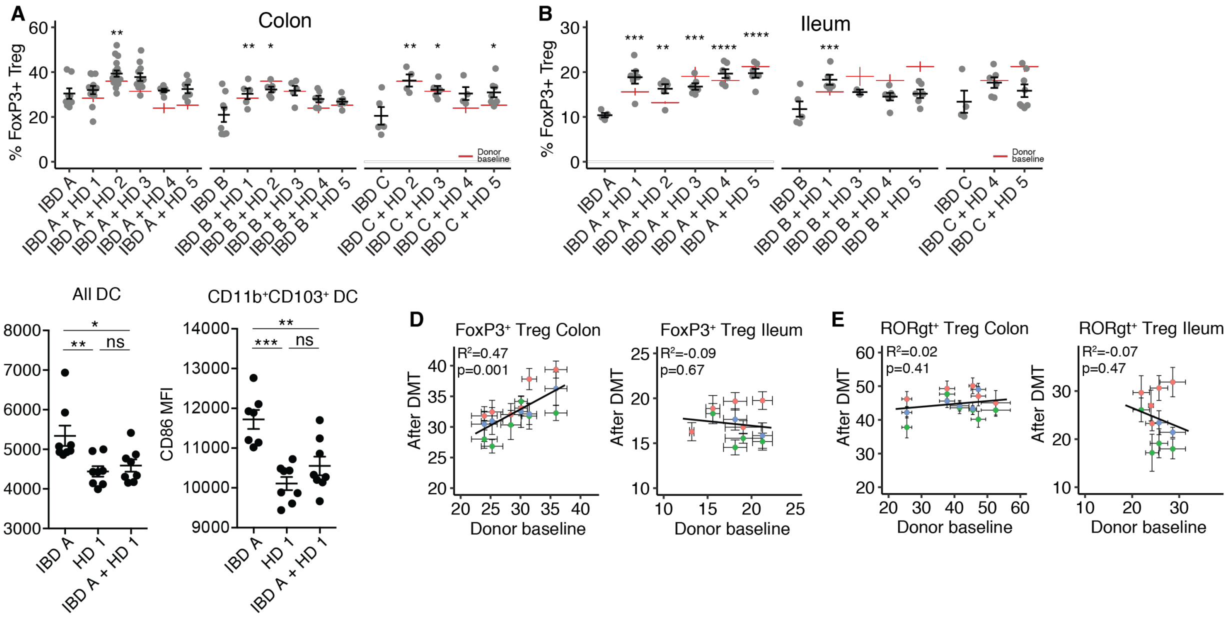


**Fig S5. Modulation of FoxP3$^+$ Treg cells and mucosal dendritic cell phenotype following microbiota transplant. (A-B)** The proportion of colon and ileum lamina propria total FoxP3$^+$ Treg cells (of live, CD4$^+$ cells) in groups of mice colonized with each IBD donor alone or three weeks following DMT with one of the five HD microbiotas. Red lines indicate the proportion of FoxP3$^+$ Treg cells induced by each HD alone. (**C**) The expression of CD86 on total dendritic cells (MHC-II$^+$, CD64$^-$, CD11c$^+$) and double-positive dendritic cells (MHC-II$^+$, CD64$^-$, CD11c$^{+,}$ CD11b$^+$, CD103$^+$) in mice colonized with donor IBD A or HD 1 alone, or following microbiota transplant with HD 1. (**D-E**) The association between the proportion of the indicated cell types in mice colonized with a HD microbiota alone, or in mice receiving the HD microbiota as a transplant following with one of three IBD microbiotas. *p<0.05, **p<0.01, ***p<0.001, ****p<0.0001, by ANOVA with Tukey correction (A-C) or f-test (E and E), ns – not significant.

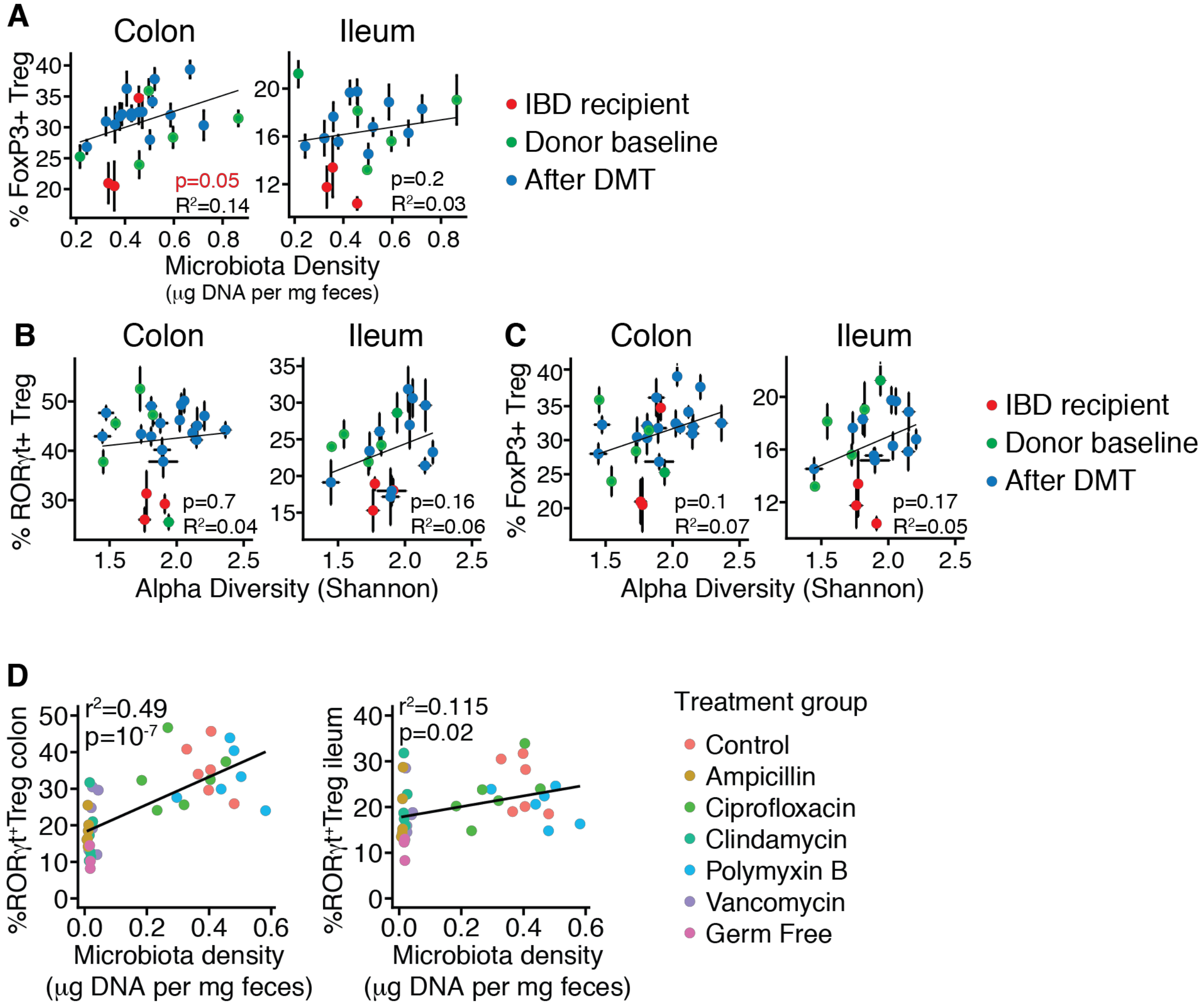


**Fig S6. The association of microbiota diversity and density with mucosal RORγt$^+$ Treg cells. (A**) The association between fecal microbiota density and the proportion of mucosal FoxP3$^+$ Treg cells in mice colonized with HD and IBD donor microbiotas alone or following DMT. (**B-C**) The association between fecal microbiota alpha diversity and the proportion of mucosal RORγt$^+$ Treg and FoxP3$^+$ Treg cells in mice colonized with HD and IBD donor microbiotas alone or following DMT. (**D**) The association between fecal microbiota density and the proportion of mucosal RORγt$^+$ Treg cells in specific pathogen free mice treated with the indicated antibiotic in drinking water. P calculated by paired t-test and in F and G by f-test.
